# Neurotrophin NT-4/5 Promotes Structural Changes in Neurons of the Developing Visual Cortex

**DOI:** 10.1101/2023.12.20.572693

**Authors:** Antonella Antonini, Sheri L. Harris, Michael P. Stryker

## Abstract

Current hypotheses on the mechanisms underlying the development and plasticity of the ocular dominance system through competitive interactions between pathways serving the two eyes strongly suggest the involvement of neurotrophins and their high affinity receptors. In the cat, infusion of the tyrosine kinase B ligand (trkB), neurotrophin-4/5 (NT-4/5), abolishes ocular dominance plasticity that follows monocular deprivation (Gillespie et al., 2000), while tyrosine kinase A and C ligands (trkA and trkC) do not have this effect. One interpretation of this finding is that NT-4/5 causes overgrowth and sprouting of thalamocortical and/or corticocortical terminals, leading to promiscuous neuronal connections which override the experience-dependent fine tuning of connections based on correlated activity. The present study tested whether neurons in cortical regions infused with NT-4/5 showed anatomical changes compatible with this hypothesis. Cats at the peak of the critical period received chronic infusion NT-4/5 into visual cortical areas 17/18 via an osmotic minipump. Visual cortical neurons were labeled in fixed slices using the DiOlistics methods (Gan et al., 2000) and analyzed in confocal microscopy. Infusion of NT-4/5 induced a significant increase of spine-like processes on primary dendrites and a distinctive sprouting of protuberances from neuronal somata in all layers. The increase of neuronal membrane was paralleled by an increase in density of the presynaptic marker synaptophysin in infused areas, suggesting an increase in the numbers of synapses. A contingent of these newly formed synapses may feed into inhibitory circuits, as suggested by an increase of GAD-65 immunostaining in NT-4/5 affected areas. These anatomical changes are consistent with the physiological changes in such animals, suggesting that excess trkB neurotrophin can stimulate the formation of promiscuous connections during the critical period.

## INTRODUCTION

In the visual cortex of mammals, the precise wiring of geniculocortical afferents and intrinsic connections that forms the organizational basis for the systems of ocular dominance (OD) and orientation selectivity develops from, and is maintained by, a fine balance of activity-based neuronal interactions (Katz and Shatz, 1996; Crair, 1999).

The anatomical and functional plastic reorganization of these systems following altered visual experience during the critical period also relies on the same activity-dependent competitive mechanisms. For example, in kittens undergoing monocular deprivation (MD) during the critical period, the weak, less active eye loses its connections and the capability to drive cortical neurons, while the connections of the more active eye are expanded along with its functional control of cortical territory (Wiesel, 1982). The central tenet of activity-dependent competition in the OD system envisions that geniculocortical afferents serving each eye compete for a limited amount of trophic messenger molecules supplied by the postsynaptic cortical neuron. In addition, the hypothesis suggests that the more active geniculocortical connections have an increased requirement for these molecules and capture them to the detriment of the less active connections, which consequently retract. A growing set of observations strongly indicates that in the visual cortex, activity-dependent competition between geniculocortical afferents involves families of neurotrophins and their high affinity tyrosine kinase (trk) receptors (Dominici, et al., 1991; Carmignoto et al., 1993; Riddle et al., 1995,1997). Indeed, if a specific neurotrophin is the limiting factor, then either local application of an excessive dose of this neurotrophin or removal of the endogenous neurotrophin should block competitive interactions among afferents.

In a physiological study using electrophysiological and optical imaging techniques in kittens during the critical period, Gillespie et al. (2000) examined the functional effects of local infusion of the trkB ligand NT-4/5 into the primary visual cortex of monocularly deprived kittens. The effects of NT-4/5 were dramatic in that OD plasticity was completely abolished—responses through the deprived eye were not lost, and the majority of cells were binocularly activated. Furthermore, orientation selectivity was lost through both eyes, although general neuronal responsiveness suffered only a mild reduction. Taken together, these observations lead to the hypothesis that large amounts of NT-4/5 prevail over competitive mechanisms driven by the difference in activity between the deprived and non deprived eyes and cause overgrowth and sprouting of thalamocortical and/or corticocortical connections. Sprouting of afferent terminals and maintenance of exuberant, nonselective connections are also consistent with the loss of orientation selectivity, a modality that requires fine tuning and specificity of circuits.

Anatomical studies are consistent with this interpretation. Local infusion of massive doses of these neurotrophins via osmotic minipumps causes the desegregation of already formed OD columns in layer 4, presumably by overgrowth of geniculate terminals (Cabelli et al., 1995,1997). Desegregation of OD columns also followed the presumed removal of endogenous neurotrophins from the visual cortex by infusion of trk-IgG fusion proteins. In addition, local infusion of another trkB ligand, brain derived neurotrophic factor (BDNF), via osmotic minipumps induces a specific sprouting in layer 4 of both deprived and non-deprived geniculate afferents in MD animals and causes sprouting of these afferents in normal animals (Hata et al., 2000). Finally, delivery of NT-4/5-coated beads into the visual cortex prevents the shrinkage of geniculate somata following MD (Riddle et al., 1995).

These anatomical studies have focused on the effects of neurotrophins on the development and plasticity of the geniculocortical afferents. In this study, we sought to explore the effect of infused NT-4/5 on the morphology of the postsynaptic neurons that receive the plethora of afferent connections.

Experimental evidence shows that neurotrophins have a preeminent role in growth and regulation of dendritic arbors of cortical neurons, with each neurotrophin eliciting both a specific pattern of dendritic changes and a modulatory effect on the action on other neurotrophins (Ruit and Snider, 1991; McAllister et al., 1995, 1997; Baker et al, 1998). However, these elegant anatomical works have focused on the early development of neurons in organotypic slices of the developing cortex in which the afferent system has been removed. In this work, we have studied neuronal morphology of an intact visual cortex infused with NT-4/5 in vivo.

Visual cortical neurons were labeled using the DiOlistics methods. We found that NT-4/5 causes in most cortical layers an excessive sprouting of spine-like processes both on dendrites and on neuronal somata.

## RESULTS

The Results are presented in three sections. In the first section, the experimental paradigm will be presented in detail. The second section describes the NT-4/5-mediated structural changes in visual cortical neurons. Finally, we analyze the pattern of immunostaining of a presynaptic marker, synaptophysin, in NT-4/5 positive and negative regions, and relate this axonal feature to the structural plasticity observed in dendrites in NT-4/5 affected areas.

### Localization of DiOlistically labeled neuronal elements relative to the infused NT-4/5

The aim of the study was to investigate possible plastic effects of neurotrophin NT-4/5 on the anatomical structure of cortical neurons. To this end, the neurotrophin was continuously infused for 4-12 days in the visual cortex of kittens during the 3rd and 4th weeks of age. No difference in the physiological effects were seen with infusions within the 4-12 day range (Gillespie et al., 2000). Cortical neurons were labeled with fluorescent dyes using the DiOlistics method (Gan et al., 2000). Fixed, 200-250μm-thick coronal sections through the lateral gyrus were shot with dye-coated particles. The particles are blown at random onto the sections and label cellular elements that come into their contact. The lipophylic dye then spreads passively along the membranes and outlines membrane elements up to their finest processes. Confocal images through successive thin planes of a labeled neuronal element reveal its anatomical features with astonishing detail (Fig. 1).

**Figure 1.**
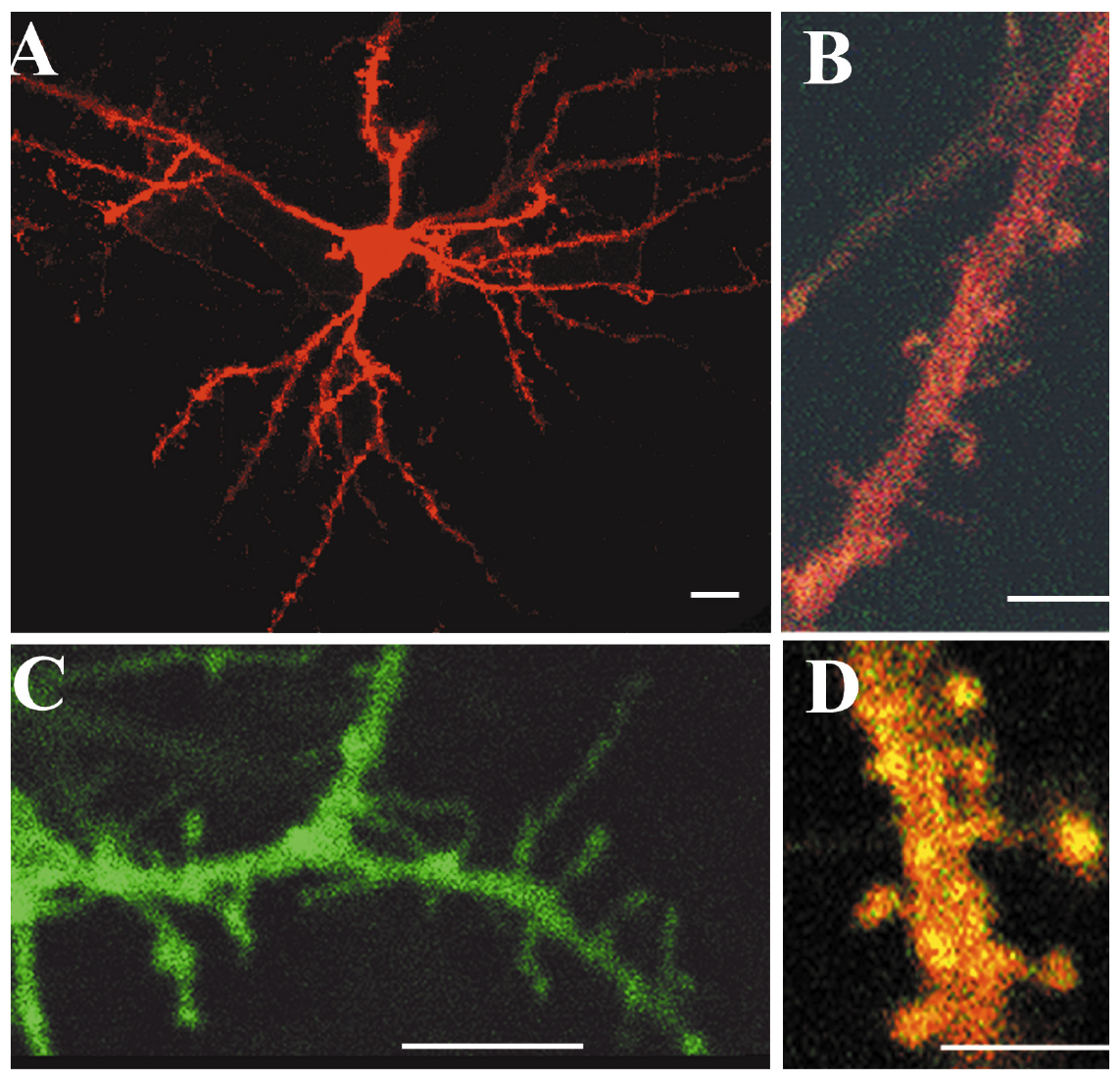
Neuronal elements of the kitten visual cortical labeled with lipophylic dyes following the DiOlistics method (Gan et al., 2000). Images were collected in laser-scanning confocal microscopy. In A, confocal images were collected as a z-stack through the depth of the section (interval between planes: 1μm), and were projected onto a single plane. In B and D, images were collected as a z-stack at an interval of 0.4μm. C was obtained with a single scan. Scales: 10μm in A and C, 5μm in B and D.

We collected neurons from the crown and the medial bank of the lateral gyrus, where areas 17 and 18 are located. Generally sections were shot with few particles to avoid a high density of labeling, particularly of axons of passage, and the consequent uncertainties in sorting out individual neurons for reconstruction. However, the use of tungsten particles coated with three different lipophylic dyes and with two by two combinations of these dyes allowed labeling neurons with a wide range of colors (Fig.1D) and facilitated the distinction between contiguous neuronal elements. Our initial plan was to serially reconstruct whole neurons in all cortical layers. However, labeled neurons were often at the very surface of the section due to the fact that tungsten particles did not penetrate deeper than 50-100μm in the tissue and clear confocal images were obtained only within 50-60μm from the top of the sample. Therefore, neurons under investigation often had cut dendrites, making complete reconstruction impossible. Qualitatively, we observed that close to the NT-4/5 infusion, the proximal segment of dendrites was unusually rich in spines. Therefore we shifted our attention to spine density measurements in both apical and basal dendrites of well-identified order.

In order to assess whether reconstructed elements (neurons or dendritic segments) were located inside or outside NT4/5 affected areas, their locations were carefully mapped in relation to blood vessels and transposed onto the adjacent 70μm-thick cortical section stained for NT-4/5 immunofluorescence. Immunopositive neuronal somata and dendrites were found in all layers. Figure 2F shows the general features of NT-4/5 immunoreactivity in cortical layers 4-6. At higher magnification it was possible to observe a dense staining of fibers (not shown). Cells in the white matter were also NT-4/5 immunopositive (Fig. 2G); these cells could have the typical pyramidal shape of subplate neurons (Fig. 2H) or could have a non-pyramidal morphology (Fig. 2I).

**Figure 2.**
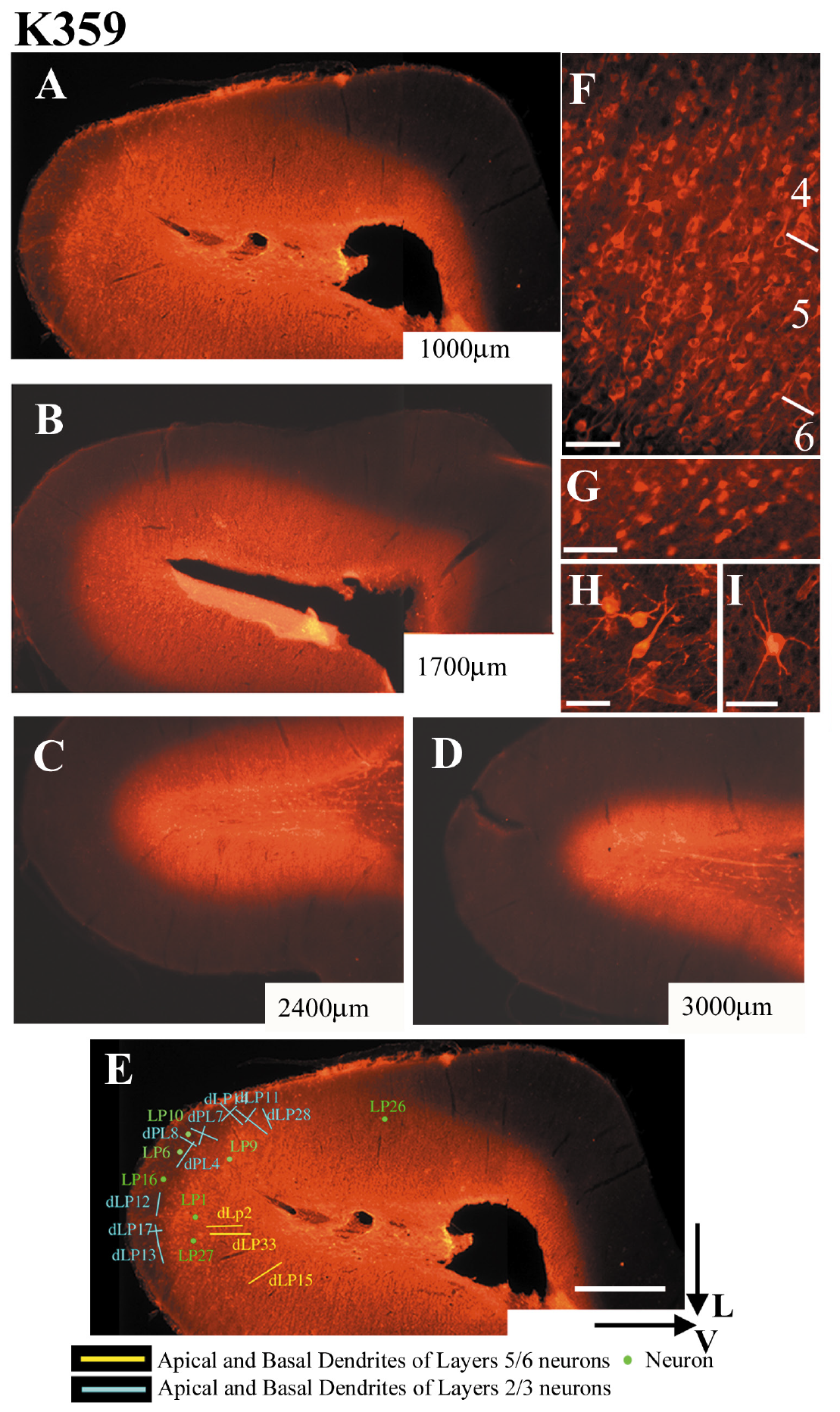
Fluorescent immunohistochemistry demonstrates the spread of the infused neurotrophin. A-D. Series of 70μm-thick coronal sections through the visual cortex infused with NT-4/5. Sections were collected at four increasing distances from the center of NT-4/5 infusion; the distances are marked at the bottom right corner of each image. Note that at 1mm from the center of the infusion, NT-4/5 is present in both the crown of the lateral gyrus and in the white matter. With increasing distance from the center of the infusion, NT-4/5 labeled area becomes restricted to the white matter. 200μm-thick sections adjacent to each one of the sections processed for NT-4/5 immunohistochemistry were used for DiOlistics experiments. In order to locate labeled neurons and dendrites relative to the NT-4/5 infused region, their position was carefully transferred onto an adjacent NT-4/5 immunostained section, as shown in E. F shows a coronal section through layer 4-6 of the visual cortex processed for fluorescent NT-4/5 immunohistochemistry. The white matter contained NT-4/5 immunopositive cells as well (G), some with pyramidal morphology resembling subplate cells (H), and others non-pyramidal (I). V and L: lateral and ventral aspects of the lateral gyrus. Scale bar for A-E: 1mm. Scale bar for F-I: 50μm.

Figure 2 (A-D) illustrates a series of 70μm thick, NT-4/5 immunostained coronal sections through the visual cortex of kitten K359 at four different levels posterior to the center of NT-4/5 infusion (see Methods). At a distance of 1000μm from the needle connected to the osmotic minipump, NT-4/5 is present throughout all cortical layers and white matter of the crown of the lateral gyrus. With increasing distance from the center of the infusion, the NT-4/5 labeled area becomes more restricted to the white matter. In Figure 2E, the position of neurons and dendrites imaged from the 200μm-thick cortical section adjacent to section A, has been transferred onto section A. In this case, all the neuronal elements are considered to be included within the NT-4/5 positive area. We considered that the reconstructed elements were located outside the neurotrophin-affected area if the adjacent section was completely NT-4/5 negative. Data obtained from these regions are considered control, since on the basis of physiological experiments on animals implanted with a NT-4/5-filled osmotic minipump, cortical regions outside the infused area have fully normal visual responses (Gillespie et al. 2000). Often, reconstructed neuronal elements were located in cortical layers free from NT-4/5 staining, while, in the same section, the underlying white matter was NT-4/5 positive. In this case neurons could have still be affected by the neurotrophin, albeit indirectly, and we classified them in a separate category (elements in border regions). The spread of NT-4/5 along the anteroposterior axis was variable. Complete labeling of cortical layers and the white matter of the entire lateral gyrus occurred within 1-2mm from the center of the infusion cannula; labeling extended in the white matter alone for another 1-1.5mm. Generally, no NT-4/5 immunostaining was found beyond 4mm from the injection site. The labeling pattern indicated that a very large expanse of area 17, 18 and possibly 19 was influenced by the infused neurotrophin.

### NT-4/5 causes proliferation of processes in visual cortical neurons

The most interesting feature found in NT-4/5 positive areas is the presence of many cortical neurons whose somata were covered with an elaborated crown of fine processes, recalling dendritic spines. Rarely were these somata found in NT-4/5 immunonegative areas: 16 out of 44 neurons (36.4%) analyzed in NT-4/5 infused areas bore processes on somata, whereas only 2 out of 28 such neurons (7.2%) were found in NT-4/5-free areas. The somatic processes could vary in shape and size: from short, stocky offshoots resembling spines to long and slender appendages similar to filopodia seen in cell cultures (Dailey and Smith, 1996). Neurons with overgrown processes could be found in cortical layers 2-6. The vast majority were pyramidal in shape; a few were also stellate cells in layer 4; however because of their small number they have not been included in the analysis.

Figure 3A illustrates examples of somata showing intricate surfaces. In contrast, Figure 3B shows the smooth membrane of normal neurons characteristic of NT-4/5 negative areas, but also present in smaller number in NT-4/5 positive areas.

**Figure 3.**
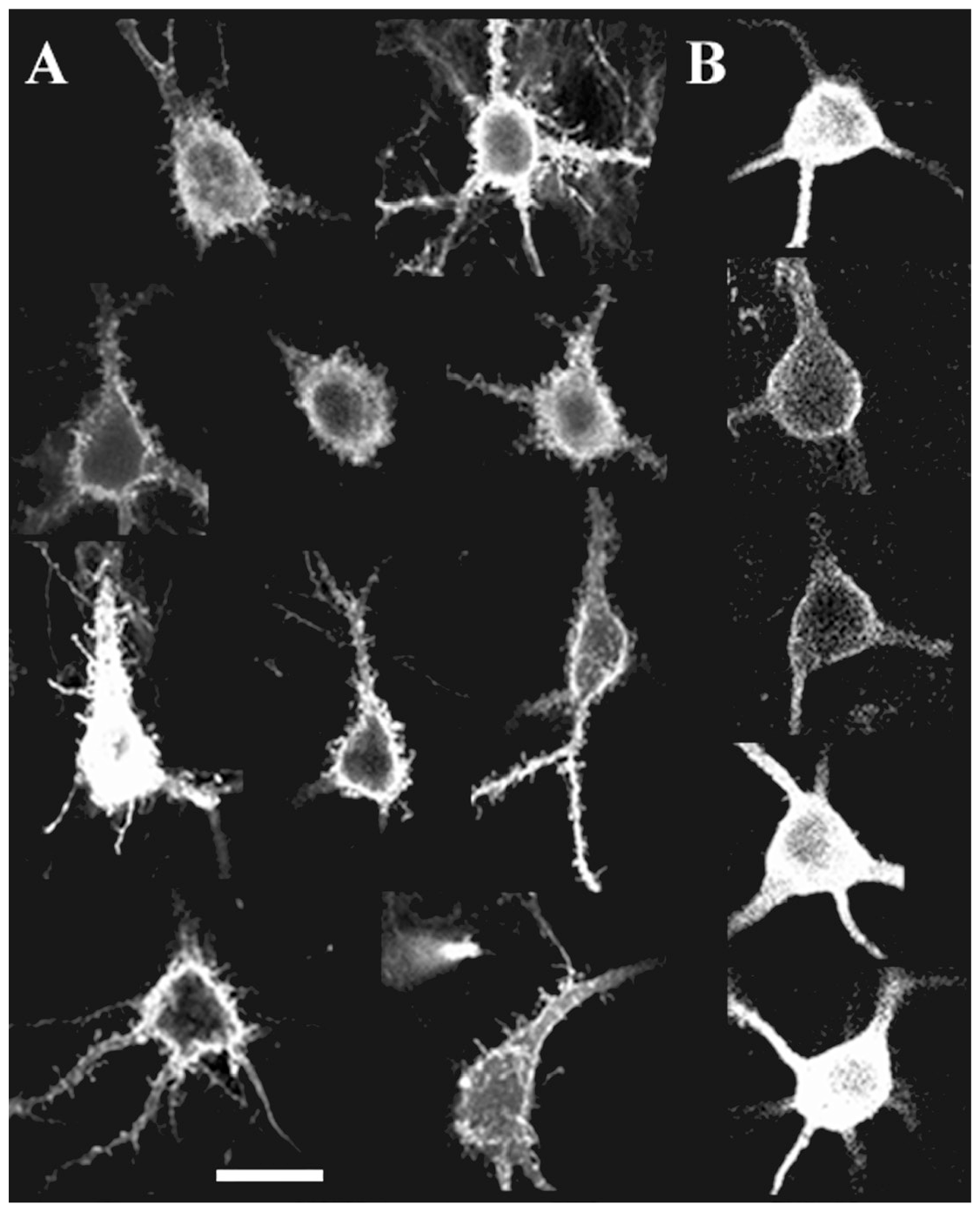
Confocal images of cortical neurons labeled using the DiOlistics method. Panels in A illustrate examples of neurons from different experiments and from different cortical layers, but they are all imaged from regions affected by NT-4/5, as established by the staining pattern in adjacent NT-4/5 immunostained sections. Note the dense sprouting of processes arising from the soma membrane. Panels in B show the smooth membrane of neurons normally found in areas both NT-4/5 positive and negative. Scale bar: 25μm

Although more conspicuous on somata, the exuberant sprouting of processes appeared to involve also dendrites. Qualitatively, we noticed very spiny dendrites in all layers, however this overgrowth was more apparent on the initial segment of apical dendrites, which are typically rather smooth.

Quantification of spine density confirmed this observation. Spines were serially mapped in confocal microscopy at 0.4 μm intervals. Spines were drawn along a portion of a dendrite and spine density was evaluated using NeuroExplorer. Dendrites were selected only if their origin (for first order dendrites) or the order of their parent branch (for second and third order dendrites) could be unmistakably identified. Data were collected mainly from neurons in layers 2/3 that are relatively small and could be easily labeled by diffusion of the lipophylic dye when a tungsten particle hit any portion of their soma or dendritic arbor. However, the amount of dye coating the tungsten particles appeared to be insufficient to diffuse throughout large membrane surfaces. Large neurons like the pyramidal cells in layers 5 and 6 were rarely labeled from the soma up to the second or third order dendrites with enough clarity to allow spine counting.

Figures 4A,B,C are plots of spine density (number of spines/μm) on apical and basal dendrites (primary, second and third order) of neurons in layers 2/3, 4 and 5/6. Neurons are divided according to their location in NT-4/5 positive areas, NT-4/5 negative areas or in border regions (see above). However for the statistical analysis (Mann-Whitney U-test) data from neurons located in NT-4/5 positive and border regions have been pooled since the two populations have similar spine counts distributions. In apical dendrites of layers 2/3 (Figure 4A, top) spine density in primary and second order dendrites was significantly higher in NT-4/5 positive areas and in border regions compared to NT-4/5 negative areas. Only the first order basal dendrites of layers 2/3 (Fig. 4A, bottom) had a higher spine density in NT-4/5 positive areas and in border regions compared to NT-4/5 negative areas. Scattergrams of Figures 4BC show that there was a strong tendency for first and second order basal and apical dendrites of neurons in layers 4 and 5/6 to bear a higher spine density in NT-4/5 infused regions, compared to NT-4/5-free regions. However, the difference between the two groups did not always reach significance. The cumulative analysis for apical and basal dendrites in all layers shows that the mean spine density is significantly higher in NT-4/5 affected regions (means: 0.18/μm outside NT-4/5 areas vs 0.39/μm inside NT-4/5 areas; p<0.0001).

**Figure 4.**
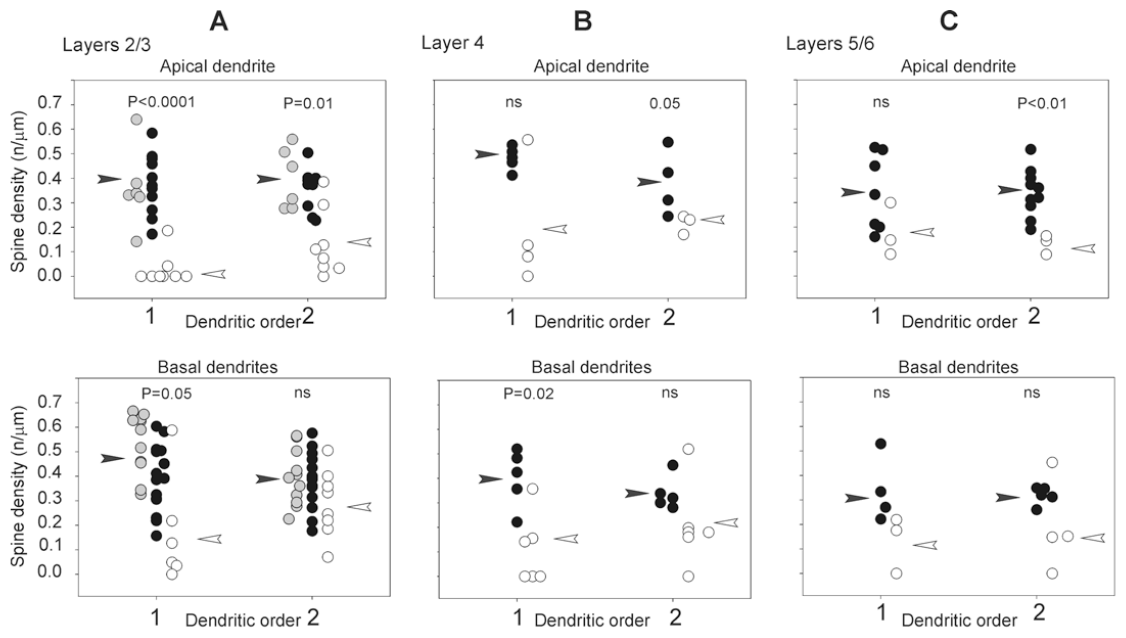
Density of spine-like processes along dendrites. Scattergram of the density of spine-like processes (ordinate: n/μm) along apical and basal dendrites of layers 2/3 (A), layer 4 (B) and layers 5/6 (C) pyramidal neurons. Processes have been counted along segments of the first and second order dendrites (abscissa). Black circles: density of processes on neurons located in a zone of areas 17 and 18 infused with NT-4/5. Empty circles: density of processes on neurons located in a NT-4/5-free areas. Gray circles: density of processes on neurons located in cortical zones in which the white matter contained NT-4/5 labeling while the gray matter did not (border zones). For the statistical comparisons (Whitney U-test) counts from the border zones have been pooled with counts in NT-4/5-infused areas. Statistical significance is shown for each comparison. ns: statistically non significant. White arrowheads indicate the mean of the distribution in NT-4/5free areas; black arrowheads indicate the mean of the distribution in NT-4/5 infused areas.

Figure 5 illustrates the percent increase in the mean density of spine-like processes between neurons inside and outside the NT-4/5-infused area. The Figure clarifies that in both apical and basal dendrites of layers 2/3 and 4 neurons, the increase in the mean density of spine-like processes induced by the NT-4/5 was higher in the first than in second order dendrites. Basal dendrites of neurons in layers 5/6 show a similar, yet mild, effect. In summary, profuse sprouting was induced by NT-4/5 infusion on the neuronal membrane of somata and on proximal portion of dendrites.

**Figure 5.**
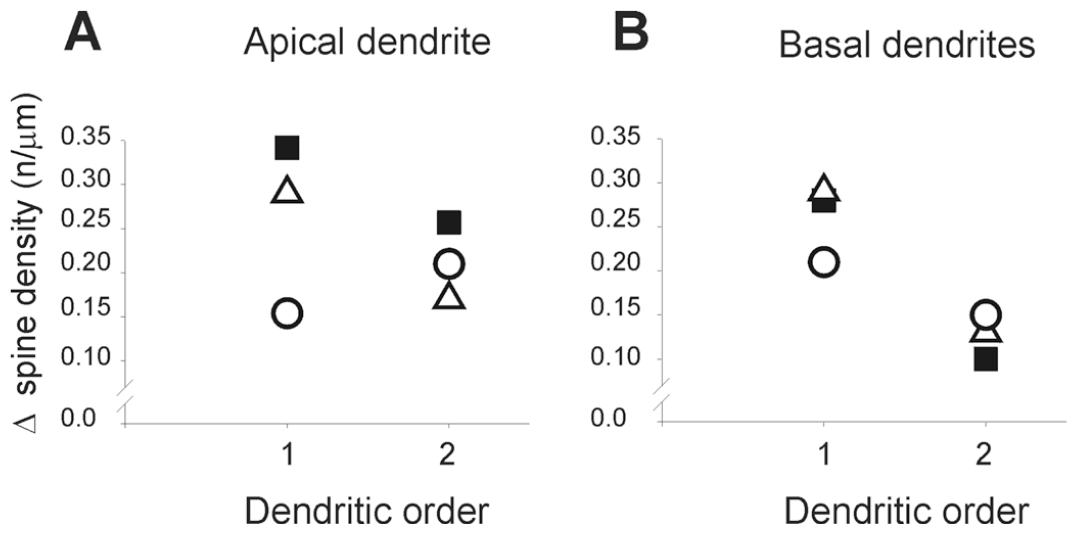
Increases of in density of spine-like processes, Percent increase in the mean density of spine-like processes on primary and second order apical (A) and basal (B) dendrites of neurons located inside or outside NT-4/5 infused region. Data for layers 2/3 are indicated with filled squares; empty triangles represent data for layer 4 and empty circles represent data for layers 5/6. Note that in layers 2/3 and 4 the largest proliferation of dendritic processes occurs for first order dendrites.

### Pattern of synaptophysin and GAD-65 immunoreactivity in NT-4/5 affected regions

An increase in dendritic protuberances resembling spines in NT-4/5 positive areas strongly suggests enhanced synaptic connections. Are the newly formed structures involved in functional synaptic contacts with presynaptic sites? If so, then presynaptic markers should increase after NT-4/5 treatment. Synaptophysin is an integral membrane glycoprotein of synaptic vesicles, and thus a marker of synaptic vesicle populations in the presynaptic terminals.

The following experiments on kitten K362 and K396 (see Methods) were aimed at studying whether local infusion of NT-4/5 in the visual cortex was accompanied by a change in synaptophysin immunoreactivity.

In confocal microscopy, synaptophysin immunofluorescence appears as distinct, bright granules present in all cortical layers. Neuronal somata, thick dendrites and blood vessels are void of immunostaining and appear as empty, black areas (Figs. 6A and 7). In some neurons, a very pale diffuse immunoprecipitate is also present in the cytoplasm (see Fig. 7). Single 2D-images of synaptophysin immunostaining were collected at random from coronal sections located along the whole antero-posterior axis of the visual cortex. Images were taken from the dorsal part of the medial bank of the lateral gyrus, where area 17 is located. In every section, special care was taken to collect the confocal image from the level along the z-axis where the immunostaining was the brightest. Images were then classified as inside or outside regions in which NT-4/5 was infused by comparison with an adjacent section stained for NT-4/5 immunoreactivity. We discarded images taken from border regions (see above). For each section we measured the average brightness of pixels over the imaged area and the average pixel brightness at 90th and 95th percentile. The density of synaptophysin immunostaining was significantly higher in visual cortical regions positive for NT-4/5 (Table 1). This evidence strongly suggests that the neurotrophin had caused a local increase in the number of synapses.

**Table 1.** Density of synaptophysin and GAD-65 immunostaining

| Table1. Density of synaptophysin and GAD-65 immunostaining |  |  |  |  |  |
| --- | --- | --- | --- | --- | --- |
| Experiment | Anti-body | area 17 | mean pixel brightness<br>(range 0-255) | 90th percentile<br>(range 0-255) | 95th percentile<br>(range 0-255) |
| K362 | Synaptophysin | inside NT-4/5 region | 68.45 | 126.64 | 150.64 |
|  |  | outside NT-4/5 region | 39.49 | 75 | 91.25 |
|  |  | Mann-Whitney U test | p<0.04 | p<0.0005 | p<0.0006 |
| K396 | Synaptophysin | inside NT-4/5 region | 107 | 191.2 | 215.6 |
|  |  | outside NT-4/5 region | 37.1 | 83 | 102.2 |
|  |  | Mann-Whitney U test | p<0.02 | p<0.02 | p<0.02 |
| K396 | GAD-65 | inside NT-4/5 region | 56.2 | 124 | 161.5 |
|  |  | outside NT-4/5 region | 30.9 | 75.2 | 100 |
|  |  | Mann-Whitney U test | p<0.003 | p<0.003 | p<0.003 |

**Figure 6.**
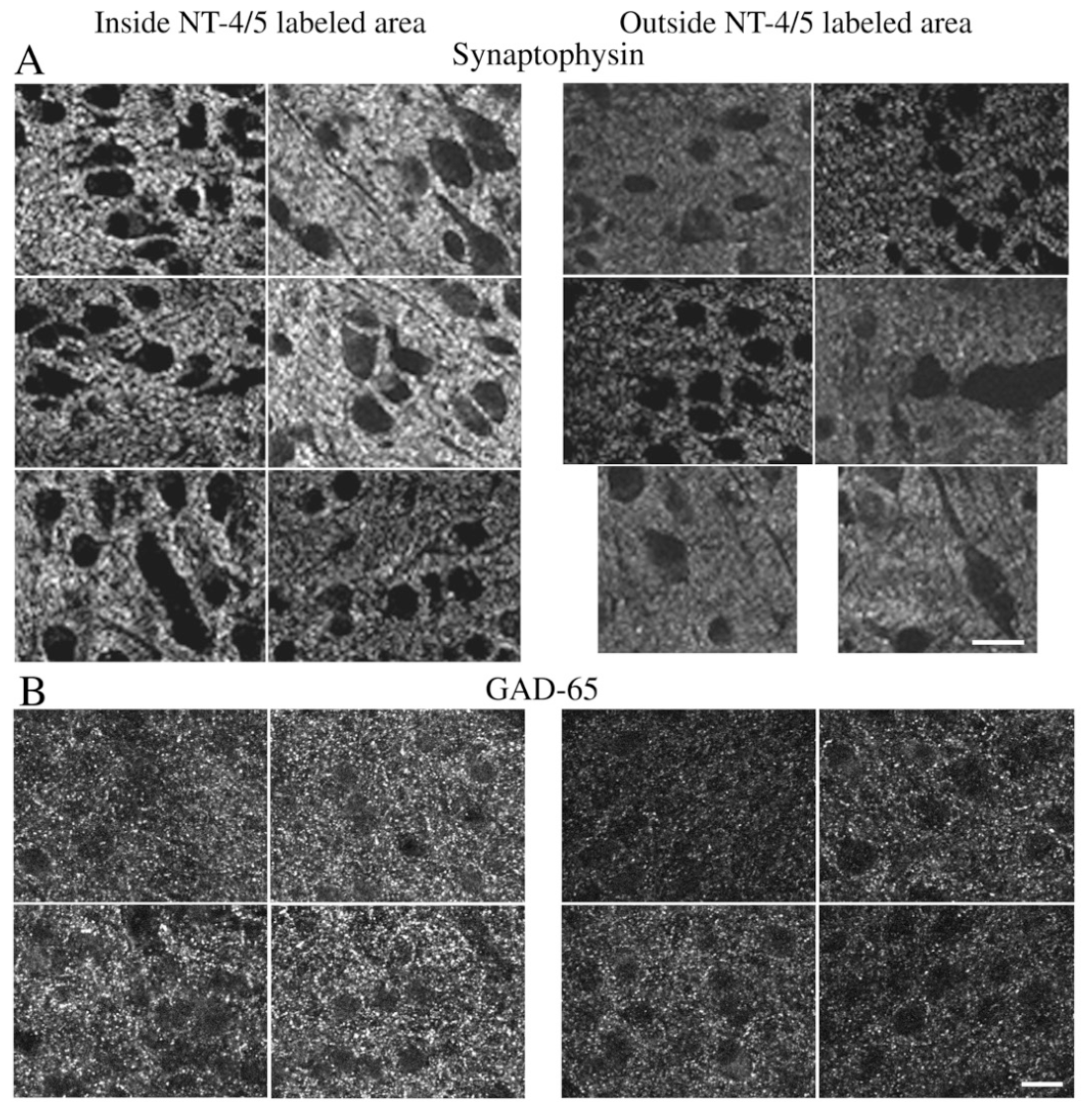
Synaptophysin and GAD-65 immunohistochemistry. Confocal mages of synaptophysin (A, case K362) and GAD-65 (B, case K396) immunostaining in regions of area 17 located inside (left panels) or outside (right panels) NT-4/5 infused region, as assessed by analyzing the spread of NT-4/5 immunoreactivity in adjacent sections. For each immunohistochemical reaction, sections have been processed in parallel and images have been collected in confocal microscopy using the same parameters of laser beam power, gain, iris and black level. The density of both synaptophysin and GAD-65 staining is significantly higher in those regions of area 17 that are affected by infused NT-4/5. Black, unstained regions are profiles of neurons, blood vessels and thick dendrites. For the determination of the density of immunostained particles of synaptophysin and GAD-65, these regions have been subtracted from the total area of each image. Scale bar 20μm.

**Figure 7.**
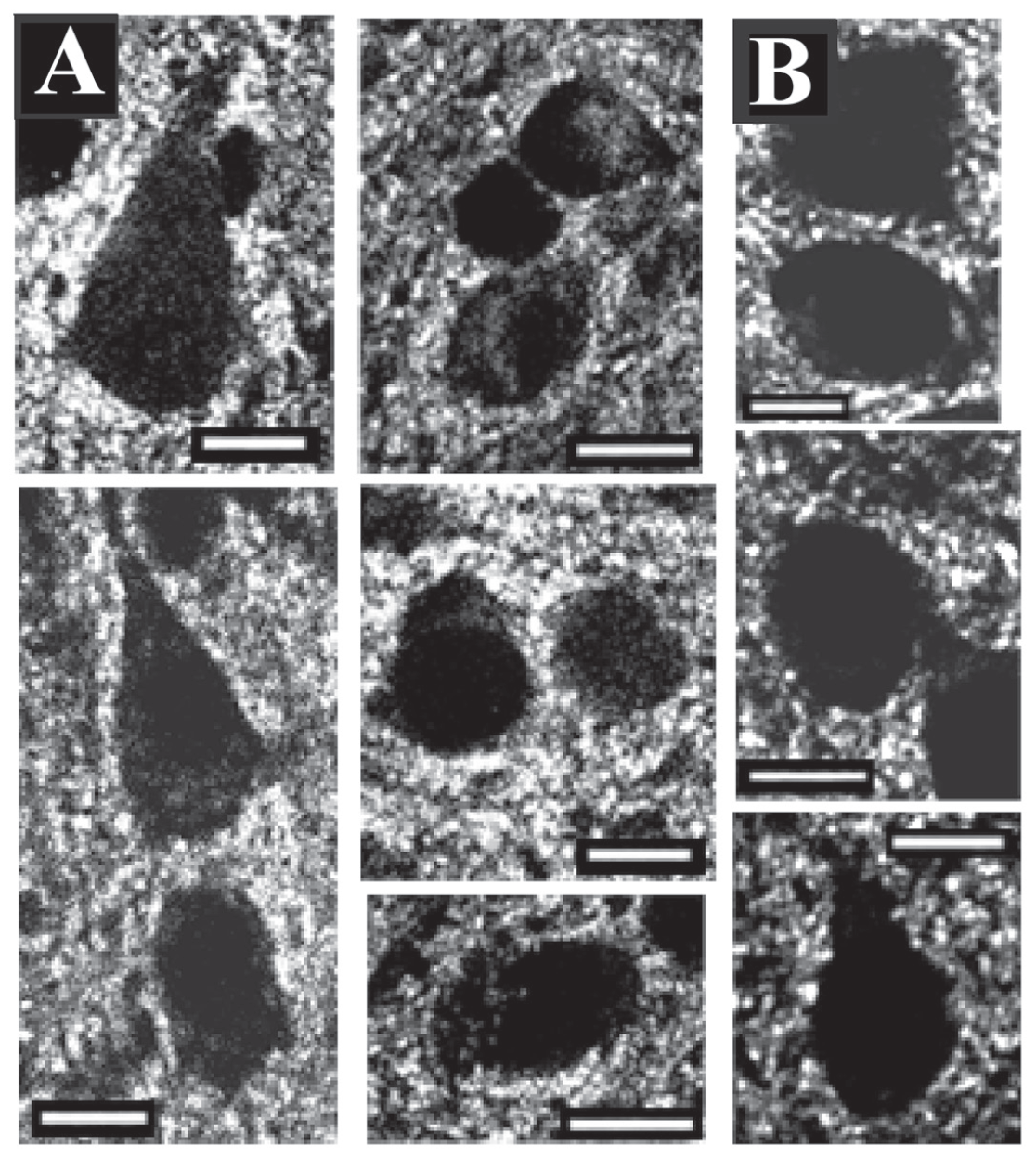
Synaptophysin granules around neuronal somata. A. In NT-4/5 infused areas neuronal somata are often surrounded by a high concentration of synaptophysin granules. B. This feature is absent in NT-4/5-free regions. Scale bar 10μm.

In regions of the visual cortex bathed by the infused NT-4/5 (i.e. regions corresponding to NT-4/5 immunopositive areas in adjacent sections), we often observed around the somata a distinct, dense crown of intensely labeled synaptophysin granules (Fig. 7). This feature was not found in NT-4/5free areas. This peculiar aspect of distribution of the presynaptic marker synaptophysin matches the localization of spine-like processes on somatic profiles and suggests that these latter might be sites of synaptic activity.

Experiments on transgenic mice overexpressing BDNF (Huang et al., 1999) or lacking trkB receptors (Rico et al., 2002) have demonstated the crucial role of this receptor in the development and maturation of inhibitory neurons. We investigated whether an excess of NT-4/5 in the developing kitten visual cortex was also accompanied by a modulation of inhibition, as measured by the density of GAD-65, a marker for inhibitory synapses. GAD-65 is an isoform of the GABA synthetic enzyme glutamic acid decarboxylase and is localized on the presynaptic terminals of GABAergic inhibitory neurons (Kaufman et al., 1991). Confocal images of GAD-65 immunostained sections were collected from the dorsal part of the lateral gyrus, where the visual cortex is located. Images were classified as inside or outside regions in which NT-4/5 was infused by comparison with an adjacent section stained for NT-4/5 immunoreactivity. Specifically, images considered outside NT-4/5 positive area were collected at a distance of 4.5mm away from NT-4/5 zone. Examples of such inside and outside fields are shown in Fig. 6B. The methods of image collection and analysis were the same as for the synaptophysin immunostaining. As shown in Table 1, the density of GAD-65 immunostaining (mean pixel brightness and brightness at 90th and 95th percentile) was significantly higher in visual cortical regions positive for NT-4/5 than in NT-4/5 free areas. It should be noted that the increase of GAD-65 immunostaining over background was still present even 1 mm beyond the NT-4/5 immunopositive area (p < 0.02 at each of the three intensity levels measured). This finding suggests that the effects of the neurotrophin extend for some distance beyond the region in which the NT-4/5 can be detected immunohistochemically.

## DISCUSSION

The present study demonstrates that excess NT-4/5 administered in vivo into the visual cortex induces a remarkable sprouting of processes from both proximal dendrites and somata of cortical neurons. The morphology of these newly formed processes closely resembles spines, although at times they appear unusually long, resembling filopodia (Figs. 1 and 3). We provide evidence that these processes might bear functional postsynaptic sites by correlating the increase in their density with the great enhancement of the presynaptic marker synaptophysin and GAD-65 at sites of NT-4/5 infusion. Such an enhancement of connectivity would be consistent with the inference from physiological studies that NT-4/5 infusion promotes promiscuous connections in developing visual cortex (Gillespie et al., 2000).

The presence of protuberances on neuronal somata has been observed in vitro (Tauer et al., 1996) and in non-mammalian species during early development (Jhaveri and Morest, 1982). In the visual cortex, we occasionally observed spine-laden somata in adult mice monocularly enucleated from P10 (Antonini, Wong and Stryker, unpublished observations).

The effect of NT-4/5 in determining new growth of dendritic and somatic processes on a cortical neuron is consistent with a retrograde action of the excess neurotrophin on the afferents to that neuron, causing growth and elaboration of their terminals. The increased presynaptic load might then increase the demand for postsynaptic sites, leading to the proliferation of spine-like processes on the postsynaptic membrane. Conversely, the effect of NT-4/5 may be explained with a direct effect of the neurotrophin on cortical neurons morphology with dendritic expansion and growth of spines (McAllister et al., 1995).

### The retrograde hypothesis

The role of NT-4/5 in the elaboration of neuronal afferents derives from the hypothesis that in the physiological condition neurotrophins be secreted by neurons on the basis of correlated activity between the neuron and its presynaptic partners. Neurotrophins would act on the presynaptic terminals as retrograde, target-derived trophic factors and would favor the stabilization and growth of the synaptic pool between the preand postsynaptic elements. Ultimately, neurotrophins’ action would be responsible for the elaboration and plasticity of presynaptic axonal terminals (Cohen-Cory and Fraser, 1995; Oppenheim, 1996; McAllister et al., 1999; Schumann, 1999).

In the mammalian visual system the retrograde action of neurotrophins has been related to the development and remodeling of eye-specific geniculocortical afferents forming OD columns, through a process involving competition (reviewed in Thoenen, 1995; Bonhoeffer, 1996; Shatz, 1997). Indeed, recent experimental evidence has suggested that in cortical layer 4 of the visual cortex, geniculocortical afferents serving each eye compete for a limited amount of a specific neurotrophin retrogradely released by the postsynaptic cortical neuron in an activity-dependent manner (Carmignoto et al., 1993; Cabelli et al., 1995; Hata et al., 2000). Several lines of evidence indicate that, in the cat, the neurotrophins of choice for this task are trkB ligands. First, in the ferret’s lateral geniculate nucleus, the neurons of origin of geniculocortical connections express both trkB and trkC receptors during development (Allendoerfer et al., 1994; Cabelli et al., 1996). Similarly, in the cat visual cortex, trkB receptors are present on geniculocortical afferents in layer 4 (Silver and Stryker, 2001). Second, maintenance and plasticity of OD columns during the critical period, appear to be overridden by an excess of BDNF and NT-4/5 or by manipulations to remove endogenous neurotrophins by infusion of chimeric trkB-IgG fusion proteins (for review see Shatz, 1997). In fact, infusion of BDNF causes the anatomical desegregation of OD columns in normal kittens (Cabelli et al., 1995; Hata et al., 2000). Desegregation of both deprived and non-deprived geniculocortical afferents with expansion of their terminal fields within layer 4 is also observed in MD animals (Hata et al., 2000). Finally, physiological experiments show that infusion of excess NT-4/5 abolishes OD plasticity after a brief period of MD (Gillespie et al., 2000). A proliferation of promiscuous connections after infusion of NT-4/5, unconfirmed by correlated activity, is also credited for the loss of orientation selectivity (Gillespie et al., 2000). Orientation selectivity relies both on the precise geometry of geniculocortical innervation (Chapman et al., 1991; Ferster et al., 1996) and on the fine tuning of intracortical circuits (Ferster and Miller, 2000), and could be disrupted by the disorganized overgrowth of connections due to an excess of neurotrophin.

Our study demonstrates that the effect of NT-4/5 is not limited to layer 4, the main geniculate recipient cortical layer.

Proliferation of dendritic and somatic processes was observed on neurons in all cellular layers. Since after infusion of BDNF, another trkB ligand, geniculate afferent sprouting is limited to layer 4 (Hata et al., 2000), the spatially diffuse action of NT-4/5 in our study suggests a target-derived growth of additional axonal terminals beyond the geniculate afferents.

Evidence from studies on BDNF indicates that inhibitory cortical circuits may be potentiated by activation of trkB receptor ligands. Indeed, studies in vitro show that application of BDNF causes differentiation of rat neocortical cultures (Nawa et al., 1993; Rutherford et al., 1997). In vivo BDNF infusion, or its overexpression in transgenic mice, has a trophic and maturational effect on Gaba-ergic neurons – both increasing soma size and neuropeptide expression (Huang et al., 1999; Nawa et al., 1994; Widmer and Hefti, 1994). Mice that overexpress BDNF also have a precocious critical period (Hanover et al., 1999), perhaps as a result of the enhancement of inhibition, since pharmacologically enhancing inhibition in normal mice can also create a precocious critical period (Fagiolini and Hensch, 2000). In our study, the preferential arrangement of newly formed spines on soma and proximal dendrites suggests that a potentiation of inhibitory circuits with sprouting of inhibitory terminals could have indeed occurred.

Plasticity of intracortical circuits may be the endpoint of a cascade of growth-inducing effects triggered by the retrograde action of the neurotrophin, or may simply derive from the direct response of cortical neurons to the trophic action of NT-4/5.

### Evidence for a direct effect of NT-4/5 on cortical neurons

McAllister et al. (1995, 1997) have shown distinct effects of different neurotrophins on growth and elaboration of dendrites in the visual cortex. These authors tested the anatomical effect of neurotrophins on neurons in layers 4, 5 and 6 in organotypic slice cultures of ferret visual cortex approximately 4 weeks earlier in development than the animals of the present study. Layer 2/3 neurons had not yet completed migration at the age tested. NT-4/5 had a widespread trophic action in all layers, although basal dendrites of layer 5 and 6 pyramidal neurons, and the apical dendrite of layers 4 and 6 pyramidal neurons responded maximally to this neurotrophin, with a dramatic increase in complexity and protospine formation. These results suggest that neurotrophins act directly on the cell through trkB receptors on its membrane rather than indirectly through elaboration of specific sets of presynaptic connections. Similar results were obtained in rat visual cortex organotypic cultures transfected with plasmids encoding for NT4/5/EGFP (enhanced green fluorescent protein). After 10 days in culture, pyramidal neurons of layers II/III and VI showed a higher spine density than controls (Wirth et al. 2003). Our data are consistent with a generalized effect of NT-4/5 on cortical neurons. Measurements from the entire neuronal population showed a dramatic increase of the mean spine density in NT-4/5-treated regions. The greatest effects were observed on the first order dendrites.

Yacoubian and Lo (2000) observed a similar pattern of specific proximal dendritic growth in organotypic cultures of ferret visual cortex overexpressing full-length trkB, whereas the truncated form of the receptor, T1, favored growth of more distal portions of the dendritic tree. The authors discussed this observation in the context of the developmental regulation of dendritic morphology, in view of the evidence that the expression ratio between the full-length trkB and T1 receptors is high in the early phases of development, when dendrites demonstrate the greatest growth, and is low later in development during the critical period, the age at which our experiments were carried out, and in adulthood (Allendoerfer et al., 1994). Although these authors focused on dendritic complexity and not on spine density, our observations of a greater effect of NT-4/5 on the proximal portions of cortical neurons, would suggest the presence of high levels of the full-length trkB isoform as if the system had reacquired some of its early developmental potential. However, evidence for up-regulation of trkB receptors by NT-4/5 are scanty: Frank et al. (1996) report that prolonged exposure to BDNF, but not NT-4/5, causes a down-regulation of trkB function.

Horch and Katz (2002) showed that the immediate neighbors of cells caused to overexpress BDNF, and the overexpressing neurons themselves by a paracrine action of BDNF, are much more dynamic, growing and losing branches and spines much more rapidly than control neurons (Horch et al., 1999). This evidence further suggests that BDNF may have a direct action on dendrites independent of presynaptic structures. Similar dynamics may also involve presynaptic partners in a developing neuronal circuit (Jontes and Smith, 2000). Part of the profuse growth induced by infused NT-4/5 has most likely occurred either through synaptic activation or by a retrograde effect of the neurotrophin on the axon, since the effect was noticeable in those layers 2/3 neurons that resided at the border of NT-4/5 diffusion (gray circles in Fig. 4A), i.e. in regions in which the soma and dendrites did not appear to be bathed in the neurotrophin, but the underlying white matter did show immunoreactivity.

### Involvement of trkB in synapse regulation

The remarkable increase of protuberances after infusion of excess NT-4/5 was accompanied by a parallel increase in the presynaptic marker synaptophysin, suggesting that the newly formed spines could bear functional synapses. These findings are consistent with in vitro and in vivo studies both on peripheral and central nervous systems showing that neurotrophins are involved in the regulation of the synaptic machinery (Snider and Lichtman, 1996; Whitford et al., 2002). Neurotrophins can alter the number of synapses (Alsina et al., 2001; Nja and Purves 1978; Rico et al., 2002), levels of synaptic vesicle proteins (Causing et al., 1997), synaptic vesicles density (Martinez et al., 1998) and both preand postsynaptic efficacy (Takei et al. 1997; Wang and Poo, 1997). It is worth mentioning that the increase of synaptophysin expression in the present experiments might not depend solely on an increase in spine density on pre-existing dendrites, but could also be dependent on new growth of spine-laden dendrites (McAllister et al. 1995, 1997; Yacoubian and Lo, 2000).

A contingent of the newly formed synaptic sites appears to be involved in inhibitory circuits, as suggested by the increase of GAD-65 immunostaining in NT-4/5 affected areas. This finding is in keeping with the evidence that trkB activation through another neurotrophin, BDNF, is able to modulate inhibitory circuitry in the visual cortex (Hensch et al., 1998; Huang et al., 1999) and the cerebellum (Rico et al., 2002).

The presence of filopodia in NT-4/5-infused cortex is a further indication that cortical neurons reacquire some immature traits, since these structures are thought to be present in early synaptogenesis. Their presence in young hippocampal cultures of postnatal brain, their high motility and the production of synapses on their tips and shafts, has led to the view that filopodia are transient precursors of more stable spines. Filopodia have been suggested to attract presynaptic terminals and guide them to the dendrite; possibly, they would form an excess of transient synaptic contacts that are then selected in an activity-dependent manner (Jontes and Smith, 2000).

### Conclusions for the role of neurotrophins in the formation of cortical circuits

The present findings provide anatomical confirmation of the suggestion made from physiological studies (Gillespie et al., 2000) that NT-4/5 infusion into critical period visual cortex stimulates the formation of promiscuous neuronal connections. One interpretation of these findings is that the mechanisms for attaining specificity in neuronal connections involve the promotion of the growth of appropriate (but not inappropriate) presynaptic connections by limiting amounts of trkB neurotrophins secreted by the postsynaptic neuron, along with the growth of appropriate postsynaptic structures. In this case, the infusion experiments would have eliminated specificity by supplying sufficient neurotrophin for all inputs, including inappropriate ones.

In the case of monocular deprivation, if limiting amounts of trkB neurotrophins are the explanation for the loss of input serving the deprived eye, then a doubling of the neurotrophin level should be sufficient to preserve inputs from both eyes. The 2-3 fold overexpression of BDNF postnatally in mouse cortex does not, however, prevent the effects of monocular deprivation and make the two eyes equally effective. Instead, such overexpression of a trkB ligand results in a precocious critical period but otherwise normal plasticity (Hanover et al., 1999). In this case, it appears that the excess neurotrophin has stimulated the precocious but otherwise normal development specifically of inhibitory neurons (Huang et al., 1999). These data then suggest that neurotrophins are not directly involved in the mechanisms of synapse specificity responsible for activity-dependent plasticity, but they are not definitive because of the possibility that compensatory mechanisms in the overexpresser mice have elevated the threshold for neurotrophin actions.

The increase of GAD-65 immunoreactivity in the areas in which NT-4/5 was infused indicates that inhibitory circuitry was also stimulated to proliferate. This finding may explain the absence of hyperexcitability or epilepsy in NT-4/5 treated cortex despite a dramatic overall increase of synaptic connections, as inferred from the increase of spine-like processes and the presynaptic marker synaptophysin, and the increases in visual responses to non optimal stimuli (Gillespie et al., 2000). The increase in inhibitory circuitry may be a direct effect of trkB stimulation. Alternatively, it may represent a homeostatic response to increased excitation, or vice versa. The time course of the physiological changes after the onset of NT-4/5 infusion does not include transient periods of hypo- or hyper-excitability, and is therefore not informative about which is more rapid or more nearly direct effect.

Modulation of inhibition has been shown to alter cortical plasticity (Hensch et al., 1998), although even dramatic increases do not prevent it, but instead reverse its direction (Reiter and Stryker, 1988; Hata and Stryker, 1994). The failure of MD to induce plasticity in visual cortex infused with NT-4/5 suggests that the increased inhibition itself is non-specific, or the combination of inhibition with an overall enhancement of synaptic activity leads to non-specific cortical circuits.

An alternative explanation for the effects of infused NT4/5 is that the high concentrations of neurotrophin cause the cortical neurons to reacquire characteristics of earlier developmental stages. In this case, the excess growth is part of a growth program, rather than part of the mechanisms that generate specificity. Such growth programs are consistent with other actions of neurotrophins in regulating neuronal size and number (Levi-Montalcini, 1966), rather than synapse specificity. Such programs might involve local synthesis of dendritic structural proteins or the rapid modulation of actin cytoskeleton dynamics (Whitford et al., 2002). These aspects of neurotrophins’ action could also be responsible for the maintenance of neuronal structure in adulthood, since complete elimination of trkB activation in the adult mouse causes loss of specific neocortical cell populations (Xu et al., 2000).

The growth-promoting effects of an excess of neurotrophins, either through the rewarding of both appropriate and inappropriate connections or through the resetting of the developmental clock to an earlier stage, can occur only during the critical period. Indeed, NT-4/5 infusion in vivo into visual cortex is without effect after the critical period and BDNF infusion causes overgrowth of geniculocortical afferents only in cortical period kittens, but not in adult cats (Hata et al., 2000). Furthermore, BDNF infusion caused a reversal of the physiological ocular dominance shift in monocularly deprived kittens but not in adult cats (Galuske et al., 1966, 2000) and, similarly, it induces synaptic potentiation of the geniculocortical pathway in young but not in adult rats (Jiang et al., 2001).

More recent studies of neurotrophin action in the cerebral cortex have turned almost exclusively to the mouse, where new genetic technology has made it possible to alter specific signaling mechanisms. In particular, the Shokat inhibitor, a chemogenetic manipulation, allows delivery of a small molecule to completely cut off the kinase activity of the trkB receptor, activation of which is the principal mechanism by which both NT-4/5 and BDNF operate (Specht and Shokat, 2002; Chen et al., 2005). Blockade of trkB kinase activity had no effect on the loss of visual response to an eye occluded during the critical period or on the increase in response to the open eye, but it completely blocked the recovery of response to the deprived eye (Kaneko et al., 2008). This finding suggested that BDNF acting on trkB was essential for the regrowth of connections serving the deprived eye, which was interpreted as the means by which responses to that eye were restored. This interpretation was reinforced by two additional findings. First, longitudinal anatomical imaging using 2-photon microscopy indicated that monocular deprivation and recovery was indeed accompanied by the loss and restoration of inputs serving the deprived eye (Sun et al., 2019). Second, measurement of the time course of production of mature BDNF revealed that it had surged by 6 hours before the recovery of visual responses to the deprived eye in both of two different circumstances in which the time to recovery differed by 24 hours (Kaneko and Stryker, 2023). These experiments, however, did not reveal the source—whether neuronal or glial—of the BDNF that mediated recovery of connections and responses. A clue to a surprising possible source was the finding that blocking the export of BDNF mRNA to the dendrites of pyramidal cells largely phenocopied the effect of blocking trkB signaling during monocular deprivation or that of an insufficiency of BDNF on the maintenance of cortical structure (Kaneko et al., 2012). Future experiments are required to determine whether BDNF infusion in the mouse has the same effects as shown here from NT-4/5 infusion in the cat.

Taken together, the results described above implicate trkB neurotrophins in the growth and development of specific types of neurons, in determining the onset of the critical period, and in the maintenance of specific populations of neurons in adult life. To date, strong evidence for their role in synapse specificity is still lacking.

## MATERIAL AND METHODS

Experiments were performed on 5 kittens (K349, K352, K359, K362 and K396) born and housed with their mother in the University of California, San Francisco cat colony. All procedures were approved by the Committee on Animal Research (University of California, San Francisco) in accordance with National Institute of Health guidelines on the Use of animals in Neuroscience Research. All efforts were made to minimize the number of animals used in this study. Neurotrophin NT-4/5 was delivered to the visual cortex through an osmotic minipump from postnatal day (P) 24 to P36, from P28 to P35, from P28 to P34, from P26 to P31 and from P25 to P29, respectively.

### Implantation of osmotic minipump

Alzet osmotic minipumps, models 2001 or 1002 (Alza, Palo Alto, CA), delivering at a rate of 1ml/hr or 0.5ml/hr respectively, were filled in sterile conditions with a solution containing 0.2mg/ml of NT-4/5 in 140mM Na-acetate, 1% Bovine Serum Albumin (BSA; Sigma, St. Louis, MO) in sodium phosphate buffer saline (PBS, 0.01M). NT-4/5 was kindly provided by Genentech Inc. (South San Francisco, CA) at a concentration of 0.6mg/ml and was diluted to match the concentration used by Gillespie et al. (2000) in a study on the physiological effects of infused NT-4/5 on the developing visual cortex. The surgical implantation of the minipumps was carried out in sterile conditions following a protocol routinely used in the laboratory. Briefly, the animal was initially anesthetized with Ketamine (10-20mg/Kg; Abbott, North Chicago, IL) supplemented with Isoflurane (1-5%) in Oxygen (1.5%) delivered through an endotracheal tube. In order to prevent gag reflexes, the insertion of the endotracheal tube was facilitated by anesthetizing the laryngeal area with 1% solution of xylocaine. The animal breathed air spontaneously; expired CO2 was monitored with a capnograph (Ohmeda, Louisville, CO). A solution of 2.5% dextrose in Lactate Ringer was delivered intravenously at a rate of 25 ml/hr. The animal was placed on a stereotaxic apparatus; the eyes were protected with an ophthalmic ointment. The minipumps were implanted bilaterally into the visual cortex. Each minipump was attached to a 30-gauge needle that was inserted in the skull through a small drilled hole and reached the visual cortex at the Horseley and Clark stereotaxic coordinates AP 0.0/-2.0 and ML 2.0. The needle was then lowered with a micromanipulator to a depth of 2mm from the surface of the brain and held in place by dental cement. The minipump itself was housed in a pocket formed in the nape of the neck. At the end of the surgery the animal was returned to the mother and had normal visual experience.

### Perfusion and Histological procedures

Four to 12 days after the minipump implantation, the animal was deeply anesthetized with an overdose of sodium thiopental (Nembutal, Abbott, 150mg/Kg), perfused transcardially with PBS followed by 2% paraformaldehyde in PBS. The brain was quickly removed from the skull and overfixed for two hours in the same fixative. A coronal cut was performed in both hemispheres in the plane of entrance of the minipump needle, dividing the hemispheres in 4 blocks: right anterior, right posterior, left anterior and left posterior. The cortical blocks were severed from subcortical structures, embedded in 4% agar and cut at the vibratome in the coronal plane. Section thickness varied according to the histological procedure and will be specified in the appropriate sections.

#### DiOlistics

Animals K349, K352 and K359 were used for this experiment. The 4 brain blocks were cut coronally in repeating series of one section 200-250 μm thick followed by two sections 70 μm thick. Sections were collected in individual wells containing 0.1M PBS and 0.05% Thimerosal (PBS-t). Each section was identified by its distance from the site of entrance of the minipump needle (i.e. from the center of NT-4/5 infusion) along the anteroposterior coordinates. The 200μm sections were used in DiOlistics experiments for neuronal labeling, while one series of 70 μm sections was used for NT-4/5 immunohistochemistry to evaluate the spread of the infused neurotrophin. This procedure allowed the assignment of neuronal elements to areas affected or non-affected by the infused NT-4/5.

In the 200-250 μm sections series cortical neurons were labeled with lipophylic dyes (DiI -1,1-dioctadecyl-3,3,3,3-tetramethylindocarbocyanine; DiO3,3-dioctadecyloxacarbocyanine perchlorate; DiD1,1 dioctadecyl 3,3,3,3 tetrametylindocarbocyanine perchlorate; Molecular Probes, Eugene, OR) of different emission wavelengths (488nm, 568nm and 647nm, respectively) using the DiOlistics procedure described by Gan et al., (2000). Briefly, 0.7 μm or 1.7 μm tungsten particles coated with one of three lipophylic dyes or with 3 different combinations of 2 lipophylic dyes (DiI+DiO, DiI+DiD, DiO+DiD) were shot by means of a helium gene gun (Helios Gene Gun System, Cat# 165-2431, Bio-Rad, CA) at 180-200psi onto cortical sections. Sections were protected from an excessive pressure wave by interposing a filter between the section and the gene gun (3μm pore size, 8x105 pores/cm^2^; Millipore). The filter also prevented large clusters of dyes from landing on the sections. No more than six sections were processed at one time and were kept in PBS-t. They were then observed and analyzed in laser-scanning confocal microscopy (Biorad, CA) by mounting them on a glass slide using an antifade mounting medium such as Vectashield (Vector, Burlingame, CA) or preferably Slow Fade (Molecular Probes, Eugene, OR) that does not contain glycerol and does not affect membrane integrity. Sections on glass slides were framed with thin strips of parafilm that provided a cushion between the slide and the coverslip and avoided deformation of the sections.

Confocal images were collected as z-stacks of 2-dimensional images (1280x1024 pixels) scrolling the depth (z-axis) of the coronal section, through the labeled element. The interval between images of the stack was 1 μm when an entire neuron was imaged at a magnification of 0.57-0.20 μm/pixel, or 0.4 μm intervals when the aim was to count dendritic spines at a magnification of 0.15-0.10 μm/pixel. The position of every neuronal element was carefully plotted and recorded onto the adjacent sections stained for NT-4/5 immunohistochemistry (see Fig. 2). At the end of the analysis, sections were transferred back to PBS-t.

Neuronal reconstruction and spine analysis were performed using the Confocal module of Neurolucida software and NeuroExplorer software (both from Microbrightfield, Colchester, VT). Statistical significance was assessed with the non parametric test Mann-Whitney U-test. Projection of 3-D stack confocal images in 2-D images was performed with Confocal Assistant software (by Todd Clark Brelje) available on-line. Anatomical figures were prepared by transferring Confocal Assistant images to PhotoShop (Adobe, San Jose, CA).

#### NT-4/5 immunohistochemistry

Immunohistochemistry for NT-4/5 was performed in all animals and started on the same day of perfusion. A series of 70 μm sections (or 100 μm sections for K362) was washed in PBS and transferred in a blocking solution containing 2.5% BSA, 3-5% Normal Horse or Normal Goat Serum and 0.3% of Triton-X in PBS-t for 2hrs to avoid non-specific labeling. Sections were then incubated for 24-48hrs in the blocking solution containing chicken anti-human NT-4/5 antibody (10 mg/ ml, Promega, Madison, WI) or rabbit anti-human NT-4/5 (1:1000, Chemicon International, Temecula, CA). The sections were subsequently rinsed 3x10min in PBS-t, incubated overnight in horse anti-chicken or goat anti-rabbit biotinylated secondary antibody (1:200; Vector Laboratories, Burlingame, CA) in blocking solution, rinsed 3x10 min in PBS-t and finally transferred to a solution of Cy3 -conjugated avidin (1:200; Jackson Immuno Research Laboratories, West Grove, PA) in PBS for 5-12 hours. Sections were analyzed in fluorescent microscopy and photographed using a digital camera (Spot software).

#### Synaptophysin immunohistochemistry

In two animals (K362 and K396), NT-4/5 was infused bilaterally for 5 days and 4 days, respectively. After perfusion, the four cortical blocks were cut at the vibratome in a series of 100 μm and 70 μm sections, respectively. One series was processed for NT-4/5 immunohistochemistry to evaluate NT-4/5 spread, as described above. Another series of sections was processed for synaptophysin fluorescent immunohistochemistry. In K362 the membrane permeant Triton-X was excluded from synaptophysin immunohistochemistry in order to combine this reaction with DiOlistics methods (data not shown). Triton-X (0.3%) was instead added to all solutions in K396). Sections were incubated in a blocking solution containing 2.5% BSA, 3% Normal donkey serum in PBS-t, then transferred for 48 hrs in a solution of mouse anti-synaptophysin (1:20, Roche Applied Science, Indianapolis, IN). Sections were then washed in PBS-t 3x10 min, incubated overnight with biotinylated anti-mouse secondary antibody (1:200, Vector Laboratories) and finally transferred to a solution of Cy2 -conjugated avidin (1:200; Jackson Immuno Research Laboratories) in PBS for 5-12 hours. After several washes in PBS-t sections were analyzed in confocal microscopy. To assess the general density of synaptophysin immunoreactivity, images were collected as single 2D-images with the filter suitable for Cy2 emission (510nm), using the same parameters of laser beam power, gain, iris and black levels for all sections. These analyses were also performed in the same day to minimize random changes in the power of the confocal microscope laser. For every section, special care was taken to collect the confocal image from the level along the z-axis where the immunostaining was the brightest. To quantify the density of synaptophysin immunostaining, we measured the average brightness of pixels from confocal images analyzed as TIFF files using IDL software (Research Systems Inc., Boulder, CO).). For each image, the area occupied by synaptophysin immunostaining was calculated by subtracting from the total area, regions occupied by somata, blood vessels and thick dendrites that appeared black. Mann-Whitney U-test was used for statistical comparisons.

#### GAD-65 immunohistochemistry

In one animal (K396) a series of 70 μm sections was processed for GAD-65 immunofluorescence according to the method described by Silver and Stryker (1999). Briefly, floating sections were incubated for 2 hours in blocking solution consisting of 0.1M PBS-t, 2.5% BSA, 0.5% Triton X-100, 3% Normal Rabbit Serum (Sigma). Sections were then incubated for 48hrs in the blocking solution containing 1:1000 rabbit anti-GAD-65 (Chemicon, Temecula, CA.). Sections were subsequently rinsed 3x10min in PBS-t, incubated overnight at 4°C in biotinylated goat anti-rabbit secondary antibody (1:200; Sigma) in blocking solution, rinsed 3x10 min in PBS-t and finally incubated overnight at 4°C. in a solution of Cy2 -conjugated avidin (1:200; Jackson Immuno Research Laboratory) in PBS-t. After a final series of three washes in PBS-t for 10 min each, sections were mounted on gelatinized microscope slides and coverslipped using as mounting medium an anti-fade solution (Vectashield; Vector). Confocal images and analysis of GAD-65 staining density were performed using the same criteria described above for synaptophysin staining density.

## ACKNOWLEDGEMENTS

Supported by NIH R37 EY02874. We thank Genentech for providing the NT-4/5 used in this study, Drs. J. Grutzendler and R. Wong for introducing us to the DiOlistics technique, and Karen MacLeod for invaluable help during the surgical procedures.

